# Model Plasma Membrane exhibits a Microemulsion in both Leaves providing a Foundation for “Rafts”

**DOI:** 10.1101/697730

**Authors:** D. W. Allender, H. Giang, M. Schick

## Abstract

We consider a model plasma membrane, one that describes the outer leaf as consisting of sphingomyelin, phosphatidylcholine, and cholesterol, and the inner leaf of phosphatidylethanolamine, phosphatidylserine, phosphatidyl-choline, and cholesterol. Their relative compositions are taken from experiment, and the cholesterol freely interchanges between leaves. Fluctuations in composition are coupled to fluctuations in the membrane height as in the Leibler-Andelman mechanism. Provided that the membrane is of relatively constant thickness, this coupling of fluctuations also provides a coupling between the composition fluctuations of the two leaves. Structure functions display, for components in both leaves, a peak at non-zero wavevector. This indicates that the disordered fluid membrane is characterized by structure on a scale given by membrane properties. From measurements on the plasma membrane, this scale is on the order of 100 nm. The theory provides a tenable basis for the origin of “rafts”.

**Statement of Significance:** The hypothesis that the plasma membrane is not homogeneous, but rather is heterogeneous, with rafts” of one composition floating in a sea of another, has overturned conventional views of this membrane and how it functions. Proteins prefer either the raft or the sea, and so are not uniformly distributed. Hence they perform more efficiently. From experiment, rafts are thought to be about 100 nm. However there is no realistic model that provides: a length scale for the rafts; a raft in both leaves of the membrane; the composition of the raft. We provide such a model. In contrast to other theories, the raft and sea are distinguished not only by composition, but also by a difference in curvature.

## 1 Introduction

The “raft” hypothesis of the mammalian plasma membrane (1, 2) has had an enormous impact on the way in which we think about this membrane. The idea posits that: both of the leaves of the membrane, rather than being compositionally uniform, are organized into regions that are relatively enriched in some phospholipids and/or cholesterol and other regions that are relatively depleted of those lipids; that in the exoplasmic leaf, the raft regions are enriched in sphingomyelin and cholesterol, while in the cytoplasmic leaf it is not yet clear in which lipids the raft region is enriched; that the regions are on the order of 100 nm; that proteins prefer one region to the other and, therefore, rather than being uniformly distributed across the membrane, are concentrated in one kind of region or the other; therefore they perform more efficiently. This encompasses the appealing idea that physical organization leads to functional organization.

There are several theories of the physical origins of rafts; they have been nicely reviewed by Schmid(3). The idea that is probably most-often cited is that rafts and the surrounding sea arise simply from liquid-liquid phase separation in the two leaves(4). This view is bolstered by the experimental observation of liquid-liquid phase separation in many model membranes consisting of at least two phospholipids and cholesterol(5). This theory, however, must contend with at least two major problems. First the idea that rafts originate from phase separation provides no explanation for the raft size, on the order of 100 nm. To address this, it has been argued that rafts arise not from phase separation that actually occurs, but rather from phase separation that would occur at a temperature lower than physiological ones; that rafts are the fluctuations associated with an unseen critical point (6). This view requires that the composition of the plasma membrane be regulated in just such a way that the system is near the critical point at such a distance that the critical fluctuations are on the order of 100 nm.

Second, and most difficult for this idea, is how to account for a raft in the cytoplasmic leaf (7), a leaf whose composition shows no tendency to undergo phase separation(8). Recall that the mechanism that drives liquid-liquid phase separation in, say, a ternary mixture is that one component, such as cholesterol, interacts more favorably with another, such as sphingomyelin, than with the third. Hence the system can lower its free energy if it phase separates into sphingomyelin-rich regions and sphingomyelin-poor ones so that the cholesterol can concentrate in the former and interact favorably with the enriched sphingomyelin there(9). As there is a large fraction of sphingomyelin in the exoplasmic leaf of the plasma membrane (10), it is not surprising that a symmetric bilayer whose leaves mimic the composition of that exoplasmic leaf undergoes phase separation(11). On the other hand, there is very little sphingomyelin in the cytoplasmic leaf of the plasma membrane. Symmetric bilayers whose leaves mimic the composition of that cytoplasmic leaf do not phase separate(8). Although phase separation in an outer leaf can induce such separation in the inner leaf if that leaf be sufficiently close to separation on its own(12, 13), the cytoplasmic leaf does not seem to be near phase separation(8). Indeed, a recent simulation showed that phase separation in the exoplasmic leaf had little effect on the properties of a model cytoplasmic leaf (14).

An alternative theory for the origin of rafts is based upon the seminal work of Leibler and Andelman (15, 16). They posited that variations of curvature in the membrane would couple to the spontaneous curvature of the lipids and thereby affect their spatial distribution. Soon after, Andelman and coworkers (17, 18) showed that this coupling could bring about modulated phases, an effect that has now been well-studied (19, 20). These phases exhibit a wavelength that is dictated by the properties of the membrane. With *σ*_*b*_ the membrane surface tension and *κ*_*b*_ its bending modulus, the characteristic length is (*κ*_*b*_*/σ*_*b*_)^1*/*2^. It was noted by one of us (21) that a modulated phase melts into a microemulsion, a fluid with structure. Like all fluids, it is characterized by a correlation length, *ξ*, but it is also characterized by a second length. This length is again the wavelenth of the modulated phase.

That “rafts” might be a manifestation of this mechanism coupling membrane curvature and compositional variations had been previously considered by Liu et al (22). However with the values of the membrane parameters they employed, they concluded that the characteristic length was much too large to describe rafts of the order of 100 nm. Ref (21), however noted that if one utilized values from Dai and Sheetz, (23), *σ*_*b*_ = 2 *×* 10^−5^*N/m* = 4.7 *×* 10^−3^*k*_*B*_*T/nm*^2^ and *κ*_*b*_ = 66*k*_*B*_*T*, then one obtains a characteristic length of 119 nm. This was cited as a reason that this mechanism could, indeed, account for the origin of rafts. Since then, microemulsions have been observed in model membranes as well as in cell-derived ones(24–26). They have not always been identified as such, however (24, 25). The experiments of ref. (26) are particularly illuminating as they show that the behavior of the characteristic length with surface tension is quite consistent with the theory (27). To observe a microemulsion in the plasma membrane with a characteristic size of 100 nm would require a neutron scattering experiment. The structure factor would have a peak at a wave vector on the order of (*σ*_*b*_*/κ*_*b*_)^1*/*2^.

There has also been much theoretical progress, including simulations of Landau modes, simulations that show what the microemulsion raft-phase might be expected to look like (3, 28, 29). The description of the plasma membrane by these models is quite simplified, utilizing only one or two order parameters. In this paper, we employ a more detailed model recently used to estimate the cholesterol distribution between leaves (30). The model, described more fully in the next section, considers the exoplasmic leaf to consist of sphingomyelin, (SM), palmitoyloleoylphosphatidylcholine, (POPC), and cholesterol, and the cytoplasmic leaf to consist of palmitoyloleoylphosphatidylethanolamine, (POPE), palmitoyloleoylphosphatidylserine, (POPS), POPC, and cholesterol. See Figure 1.

**Figure 1:**
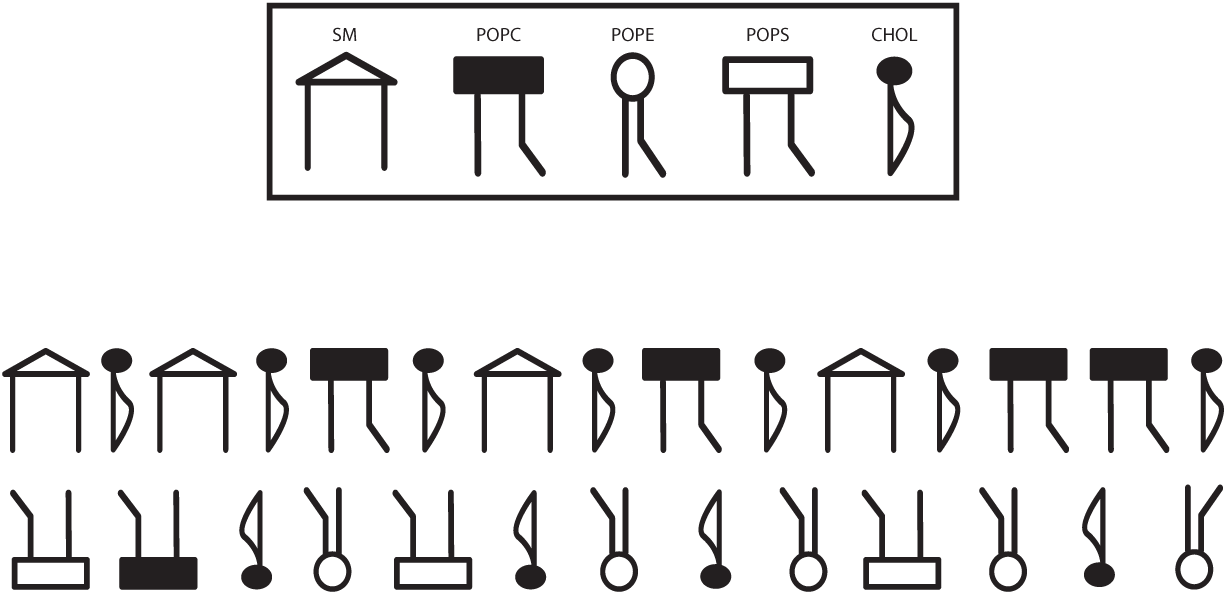
Schematic of the model plasma membrane.

We calculate the free energy of the system up to, and including, Gaussian fluctuations. From this we obtain all twenty-eight structure functions. Several of them, in both leaves, show a peak at a wavevector on the order of (*σ/κ*)^1*/*2^ characteristic of a microemulsion and, in particular, of a raft on the order of 100nm. From these structure functions, a picture of a raft emerges as a region enriched, in the outer leaf, with SM and cholesterol and enriched in the inner leaf with POPE and POPC, floating in a sea which is POPC-rich in the outer leaf, and rich in POPS and cholesterol in the inner leaf. The raft and the sea are distinguished, in both leaves, by a difference in curvature.

## 2 Theoretical Model

We begin with a model of the plasma membrane utilized recently (30). The exoplasmic leaf consists of 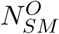 molecules of SM, 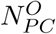 molecules of POPC, and 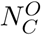 molecules of cholesterol; the cytoplasmic leaf consists of 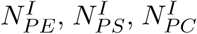, and 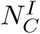 molecules of POPE, POPS, POPC, and cholesterol respectively. It is convenient to define the mol fractions in each leaf

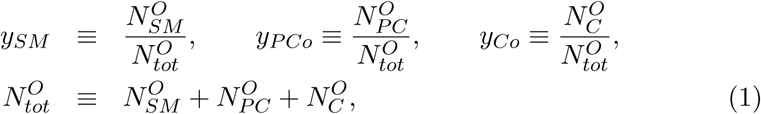

and

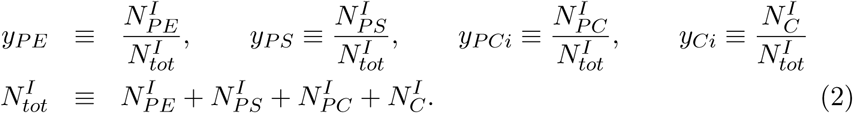

We work in an ensemble in which the areas of the leaves are fixed, and take these areas to be equal. Given that the fractional difference in areas of the two leaves is on the order of the ratio of the thickness of the plasma membrane to the size of the cell, about 10^−3^, this assumption is reasonable. We further assume that the leaves are incompressible so that the area, *A*, of the leaves is determined by their composition. Denoting the area per phospholipid by *a* = 0.7nm^2^(31) and that per cholesterol by *r* × *a* = 0.42 nm^2^ (32), with *r* = 0.6, we can write the area of each leaf in terms of the number of molecules in the outer leaf

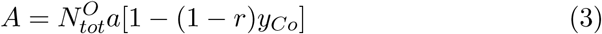

or the inner leaf

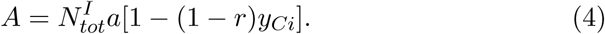

Thus the number densities of particles in the two leaves are

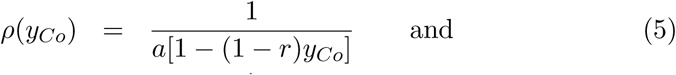

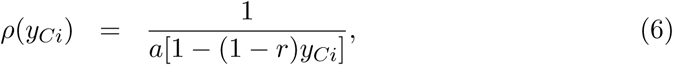

respectively.

We assume that the components in the different leaves do not interact explicitly with one another. As a consequence the free energy can be written

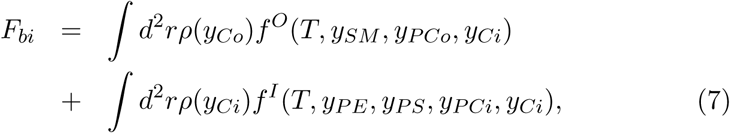

where *f* ^*O*^ and *f* ^*I*^ are free energies per particle, and *T* is the temperature.

The components within each leaf inteact pairwise. In the simplest approximation, mean-field, or regular solution, theory the free energies per particle are given by interaction and entropy terms

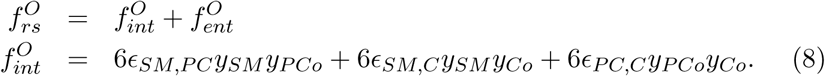

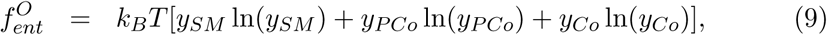

and

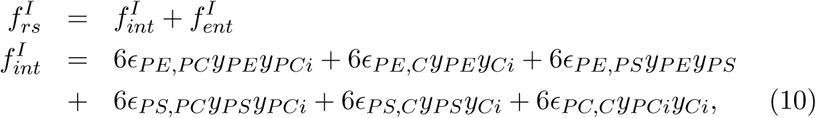

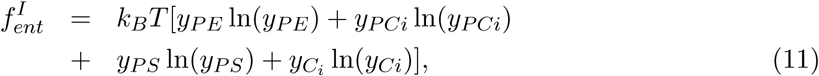

Terms linear in the component mol fractions have been ignored as they only affect the absolute values of the component chemical potentials. The epsilons in the above are interaction energies which will be given later.

We now consider the bending energy that must be overcome to bring the two monolayers together into a bilayer. The bending energy of the two leaves can be written in the Monge representation as

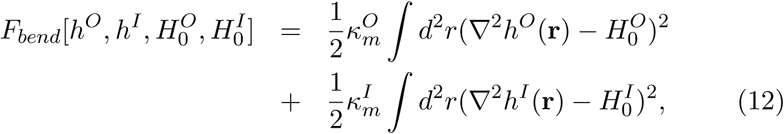

where 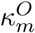 and 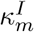 are the bending modulii of the monolayers, 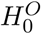 and 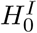 are their spontaneous curvatures, and *h*^*O*^(**r**) and *h*^*I*^(**r**) are the heights of the leaves measured from each point **r** in some external plane. We first assume that the bilayer is flat, so that ∇^2^*h*^*I*^ = ∇^2^*h*^*O*^ = 0. The bending free energy reduces to

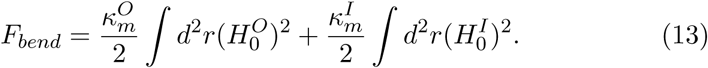

Thus the total free energy is now

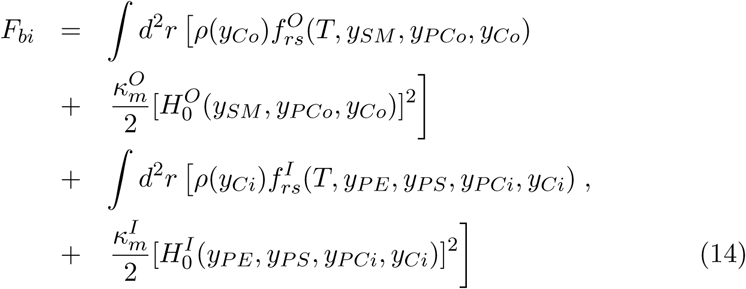

We must still specify the interaction energies and the spontaneous curvatures of the two leaves. We use the same interaction energies as previously, (30). The interactions which involve cholesterol are: *ϵ*_*SM,C*_ */k*_*B*_*T* = −0.58 *ϵ*_*PE,C*_ */k*_*B*_*T* = 0.28, *ϵ*_*PC,C*_ */k*_*B*_*T* = 0.20, and *ϵ*_*PS,C*_ */k*_*B*_*T* = −.06. The interaction between phospholipids are weaker and we take them all to vanish; i.e. *ϵ*_*SM,PC*_ = *ϵ*_*PE,PS*_ = *ϵ*_*PE,PC*_ = *ϵ*_*PS,PC*_ = 0.

As to the spontaneous curvatures, a word must be said about the sign convention. As noted earlier, the displacements of the leaflets above a position **r** on an external plane are denoted *h*^*O*^(**r**) and *h*^*I*^(**r**). If *h*^*O*^ and *h*^*I*^ are arbitrarily taken to increase in the direction from the exoplasmic leaf to the cytoplasmic leaf, then a lipid with small headgroup and wide tails has a spontaneous curvature that is negative if it is on the exoplasmic leaf. The curvature of the same lipid is positive if it is on the cytoplasmic leaf. The product of the leaflets’ spontaneous curvatures and their bending moduli was obtained from simulation in ref.(30). They were found to depend on the mol fraction of cholesterol in the leaf. In the relevant range of cholesterol fraction, it was determined that in the exoplasmic leaf, the product of monolayer bending modulus and spontaneous curvature was well fit by the form

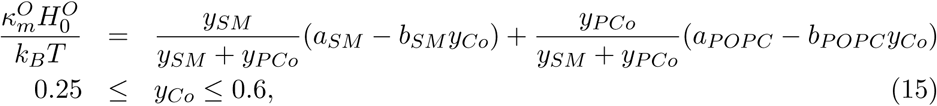

where, in units of nm^−1^, *a*_*SM*_ = 6.438, *b*_*SM*_ = 18.58, *a*_*POP C*_ = 1.341, and *b*_*POP C*_ = 9.267. Note that the contribution of cholesterol to the leaflet’s spontaneous curvature is only through its effect on the phospholipids (33); it tends to enlarge the area taken up by their acyl chains (30).

A similar fit to the product of the monolayer bending modulus and spontaneous curvature of the cytoplasmic leaf was found to be

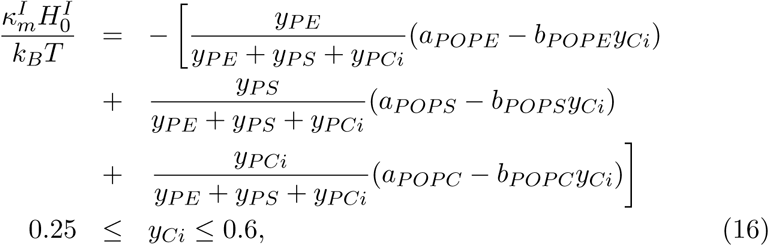

with *a*_*POP E*_ = −3.156, *b*_*POP E*_ = 2.71, and *a*_*POP S*_ = −1.223, *b*_*POP S*_ = 3.35, again, all in nm^−1^.

Lastly we assume that the bending moduli of the two leaves are equal, and equal to one half of the bending modulus of the bilayer,

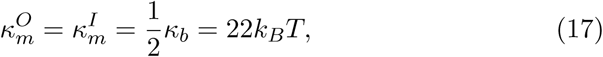

where we have used the bending modulus, *κ*_*b*_, of the membrane of red blood cells(34).

The description of the free energy of a non-fluctuating model membrane is now complete, but the mol fractions of the various components that describe the mammalian plasma membrane have not yet been specified. To do so, we begin by noting that the free energy per particle, *f* ^*O*^, of the outer leaf is a function of the mol fractions *y*_*SM*_, *y*_*PCo*_, *y*_*Co*_, and the free energy per particle of the inner leaf, *f* ^*I*^, is a function of *y*_*PE*_, *y*_*PS*_, *y*_*PCi*_, and *y*_*Ci*_. However as

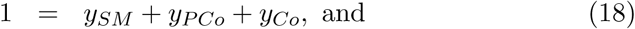

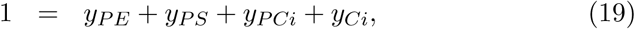

only five of the seven mol fractions are independent. Furthermore, as cholesterol can flip-flop between leaves rapidly (35–37), the chemical potentials of the cholesterol in the two leaves should be equal:

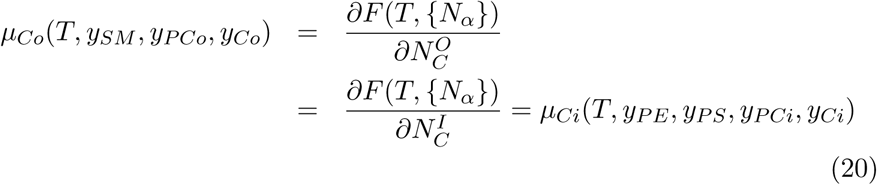

We further restrict the state of the system to be such that the mol fraction of the cholesterol in the bilayer, *x*_*c*_, be equal to 0.4 (38). Utilizing the equality of the areas of the two leaves as expressed in Eqns 3 and 4, we can write

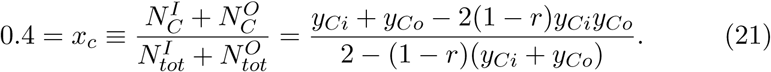

Specification of the system is completed by requiring that the mol fractions of SM and PC in the outer leaf be in the ratio 1:1, and the ratios of the mol fractions of PE to PS to PC be 5:3:1 (10).

Given these requirements, ref.(30) finds that the equilibrium mol fractions of the various components, denoted here by an overbar, are, in the outer leaf

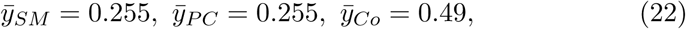

and in the inner leaf

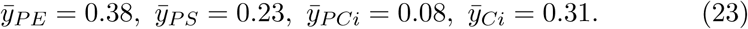

The per cent of the total cholesterol that is in the inner leaf is 36.5%.

This completes the review of the model plasma membrane utilized earlier (30). In this paper we extend this model free energy to include Gaussian fluctuations so that we can obtain the structure functions that the model predicts. These structure functions are, of course, quite informative. We show that they do provide some insight into the nature of “rafts”.

To extend the free energy of Eq.(14), we define the fluctuation of the mol fraction of the *ν* component by

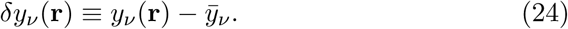

Because the mol fractions now fluctuate about their average, we must add to the free energy the usual gradient squared penalty for large fluctuations;

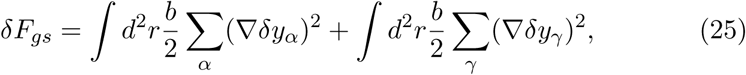

with *α* taking the values of the outer leaf, SM, PCo, and Co, and *γ* the values of the inner leaf PE, PS, PCi, and Ci. We take (39) the energy cost *b* = 5*k*_*B*_*T*.

The contribution to the free energy, Eq. (14), from the regular solution theory is expanded to second order to give the additional term

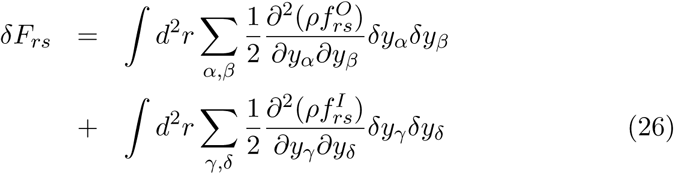

Similarly the terms in Eq. 14 which are quadratic in the spontaneous curvatures contribute

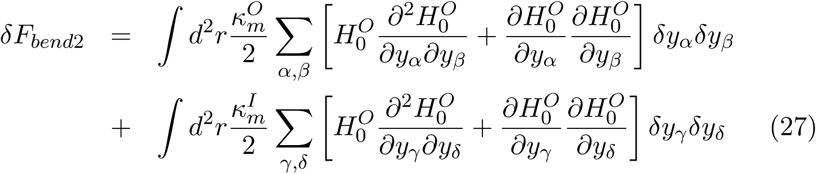

Because the membrane’s spontaneous curvature can now vary locally, we must permit the membrane’s local curvatures ∇^2^*h*^*O*^ and ∇^2^*h*^*I*^ to vary from being flat. Therefore the bending energy of Eq.(12) contributes

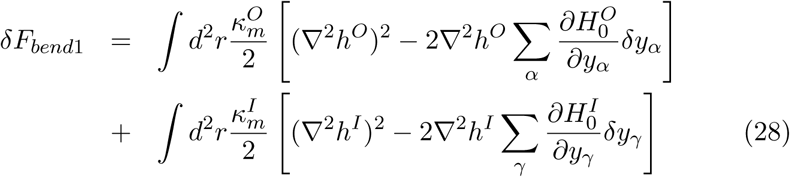

Finally, because the membrane is no longer flat, there is a surface free energy

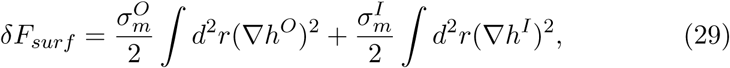

where 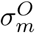 and 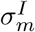 are the surface tensions of the two leaves. The total contribution of Gaussian fluctuations to the free energy is

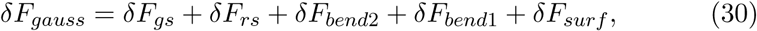

where the five terms are given in Eqs. (25-29)

It is convenient at this point to express all fluctuations in terms of the Fourier transforms

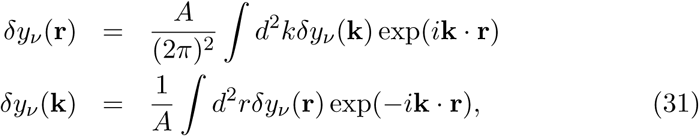

and similarly for *h*(**k**). We shall also assume that

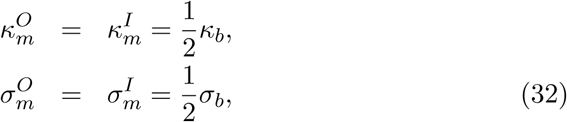

where we shall take *κ*_*b*_ = 44*k*_*B*_*T* (34) for the bilayer bending modulus and *σ*_*b*_ = 4.7 *×* 10^−3^*k*_*B*_*T* /nm^2^ (23). Note that the only terms which involve the fluctuations of the membrane height are *δF*_*bend*1_ and *δF*_*surf*_:

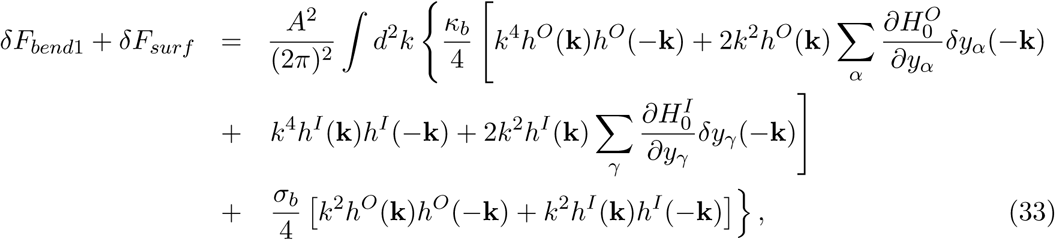

where terms linear in the fluctuations have integrated to zero;

We now assume that the thickness of the plasma membrane is constant, which implies that

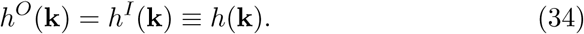

With this assumption, the above equation simplifies to

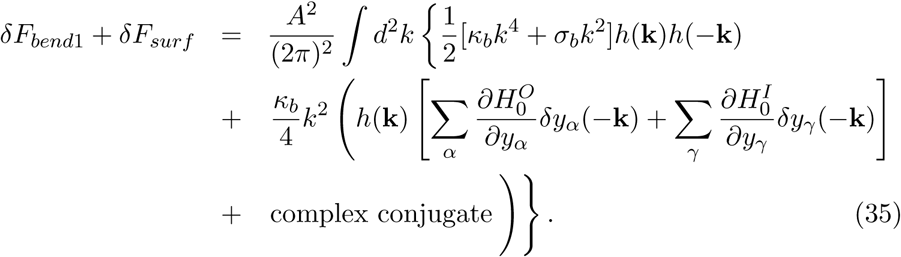

Because we are not interested here in the actual measurement of the fluctuations of the membrane height, we eliminate them by minimizing the above contribution to the free energy with respect to *h*(**k**). We obtain

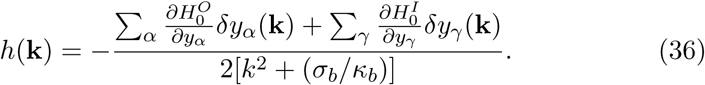

This result shows that, at large wavevectors, or short distances, (*k*^2^ *>> σ*_*b*_*/κ*_*b*_), the local membrane curvature −*k*^2^*h*(**k**), approaches the change in the membrane spontaneous curvature resulting from the composition fluctuations. However at larger distances the response of the membrane to changes in the spontaneous curvature is suppressed by the surface tension. The characteristic length below which the membrane can respond to changes in the spontaneous curvature is (*κ*_*b*_*/σ*_*b*_)^1*/*2^. With the values of *κ*_*b*_ = 44*k*_*B*_*T* (34) and *σ*_*b*_ = 4.7*×*10^−3^*k*_*B*_*T* /nm^2^ (23), this maximum length is 97 nm. This provides an explanation for the characteristic size of rafts, and is strong evidence that their origin lies in the coupling of the local curvature of the plasma membrane to its local spontaneous curvature, as suggested previously (21).

We now substitute this result into the free energy contributions of Eq. (33). Noting that these contributions contain a term *h*(**k**)*h*(−**k**), we see that it will contain a term proportional to

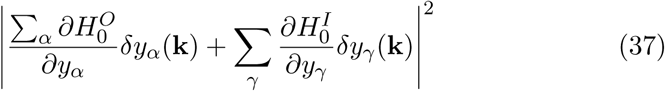

which couples composition fluctuations in the inner and outer leaves. Thus the forces that tend to keep constant the thickness of the plasma membrane, such as those of the hydrophobic effect, provide a means to couple the compositional fluctuations, i.e. the rafts, of one leaf and the other (27).

We next express all the other second-order contributions to the free energy, Eq. (30), in terms of the Fourier components *δy*_*ν*_ (**k**). Lastly, we recall that only five of the seven composition fluctuations are independent, as expressed in Eq. (18). This implies that

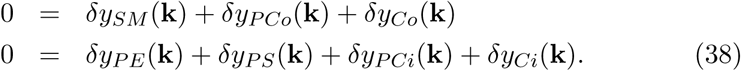

We therefore eliminate *δy*_*PCo*_(**k**) and *δy*_*PCi*_(**k**) in terns of the other fluctuations. We finally arrive at an expression for the Gaussian fluctuation contribution to the free energy. It can be written compactly as

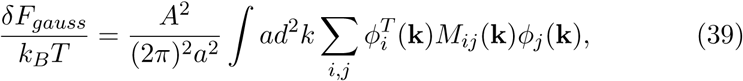

where 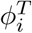 is the row matrix

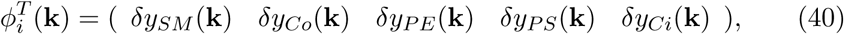

*ϕ*_*j*_ the corresponding column matrix, and *M*_*ij*_(**k**) is a 5*×*5 symmetric matrix. Its fifteen independent elements are given in the Supplementary Information.

The desired structure functions are now obtained from

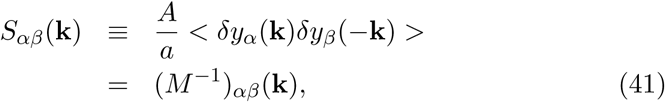

where the brackets denote an ensemble average. The structure factors so obtained are a function only of the magnitude, *k*, of the wavevector, and not of its direction: *S*(**k**) = *S*(*k*).

## 3 Results

We show in the next several figures results for some of the twenty-eight correlation functions, of which fifteen are independent. Given the large magnitude of the spontaneous curvature of PE due to its small headgroup, one would expect the effect of the coupling of membrane curvature to spontaneous curvature to be large for this molecule. Fig. 2 shows *S*_*PE,P E*_ (*q*), where it is plotted against the wavevector *k* in units of the wavevector (*σ*_*b*_*/κ*_*b*_)^1*/*2^ which arises from the coupling of concentration and spontaneous curvature; *q* ≡ *k*(*κ*_*b*_*/σ*_*b*_)^1*/*2^. This structure function clearly shows that the inner leaflet is a microemulsion because the peak of the structure function is at a non-zero wavevector which indicates that the disordered fluid nevertheless has structure at a certain wavelength *λ* = (2*π/q*)(*κ*_*b*_*/σ*)^1*/*2^. As the peak in this structure factor is at a *q* value of about eight, the corresponding wavelength is 76 nm. The same behavior is seen in the structure function *S*_*Ci,Ci*_ of the cholesterol in the inner leaf, as shown in Fig. 3. From the cross-structure function, *S*_*PE,Ci*_, not shown, we find that the PE and the cholesterol in the inner leaf are anti-correlated. From other cross-stucture functions of the inner leaf, we find that the PC tends to aggregate with the PE, and the PS with the cholesterol.

**Figure 2:**
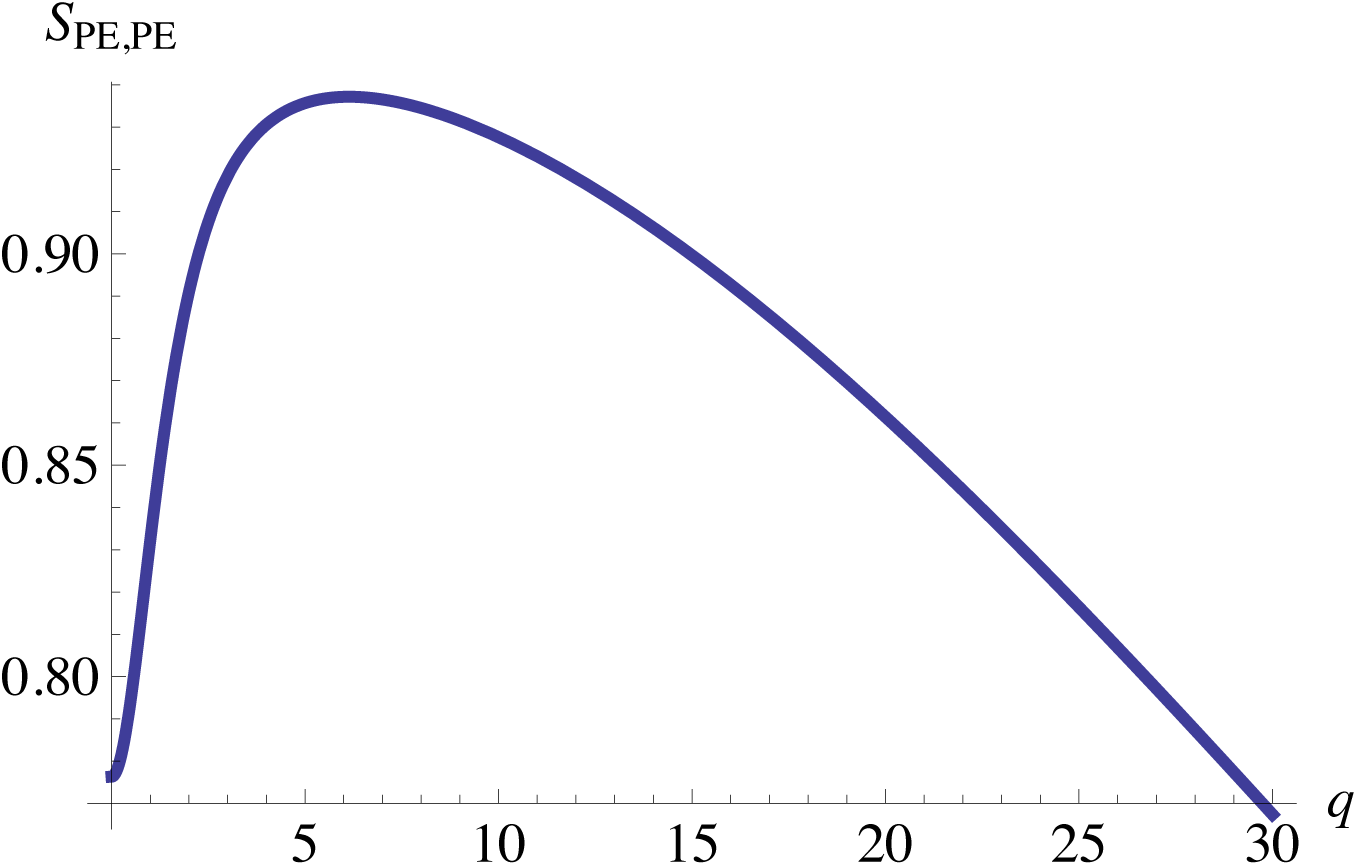
PE-PE structure function plotted versus the dimensionless wavevector *q* ≡ *k*(*κ*_*b*_*/σ*_*b*_)^1*/*2^. The structure function at zero *q* is *S*_*PE,P E*_ (0) = 0.78.

**Figure 3:**
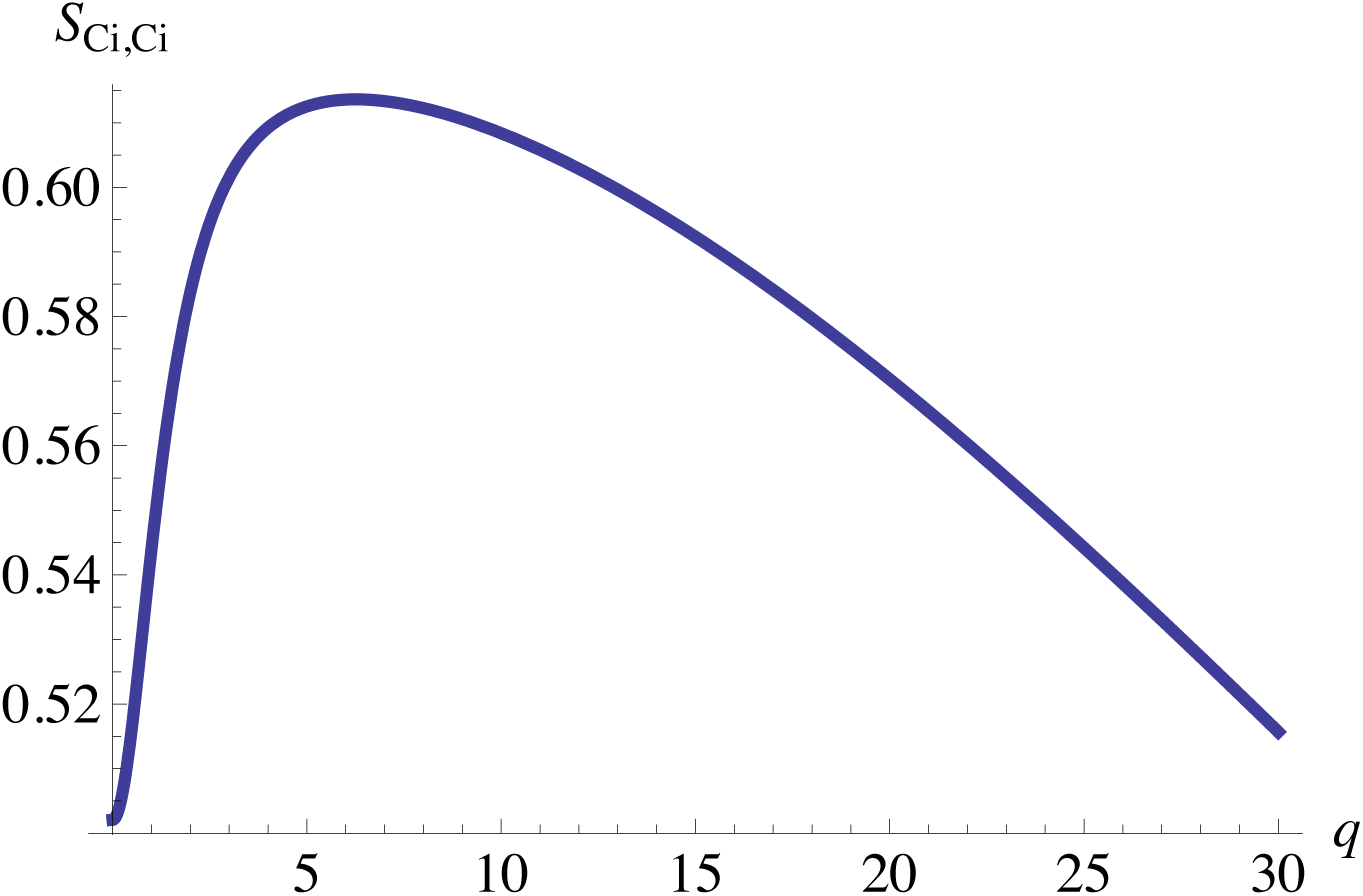
Structure function of the cholesterol in the inner leaf plotted verus the dimensionless wavevector *q* ≡ *k*(*κ*_*b*_*/σ*_*b*_)^1*/*2^. The structure function at zero *q* is *S*_*Ci,Ci*_(0) = 0.50.

Due to the coupling between leaves which, as discussed above, occurs quite simply within this theory, the microemulsion behavior seen in the inner leaf is also expressed in the outer leaf. This can be seen in the cholesterol-cholesterol structure function of the outer leaf shown in Fig. 4. We note that the value at the peak in *S*_*Co,Co*_ is nearly twice the value at *q* = 0. The structure functions *S*_*SM,SM*_ and *S*_*PCo,PCo*_ are similar. The cross-structure functions between SM and cholesterol shows that their fluctuations are correlated, while the structure function between SM and PC indicates that these components are anti-correlated. Thus the raft in the outer leaf consists of regions of enriched SM and cholesterol floating in a sea of enriched PC.

**Figure 4:**
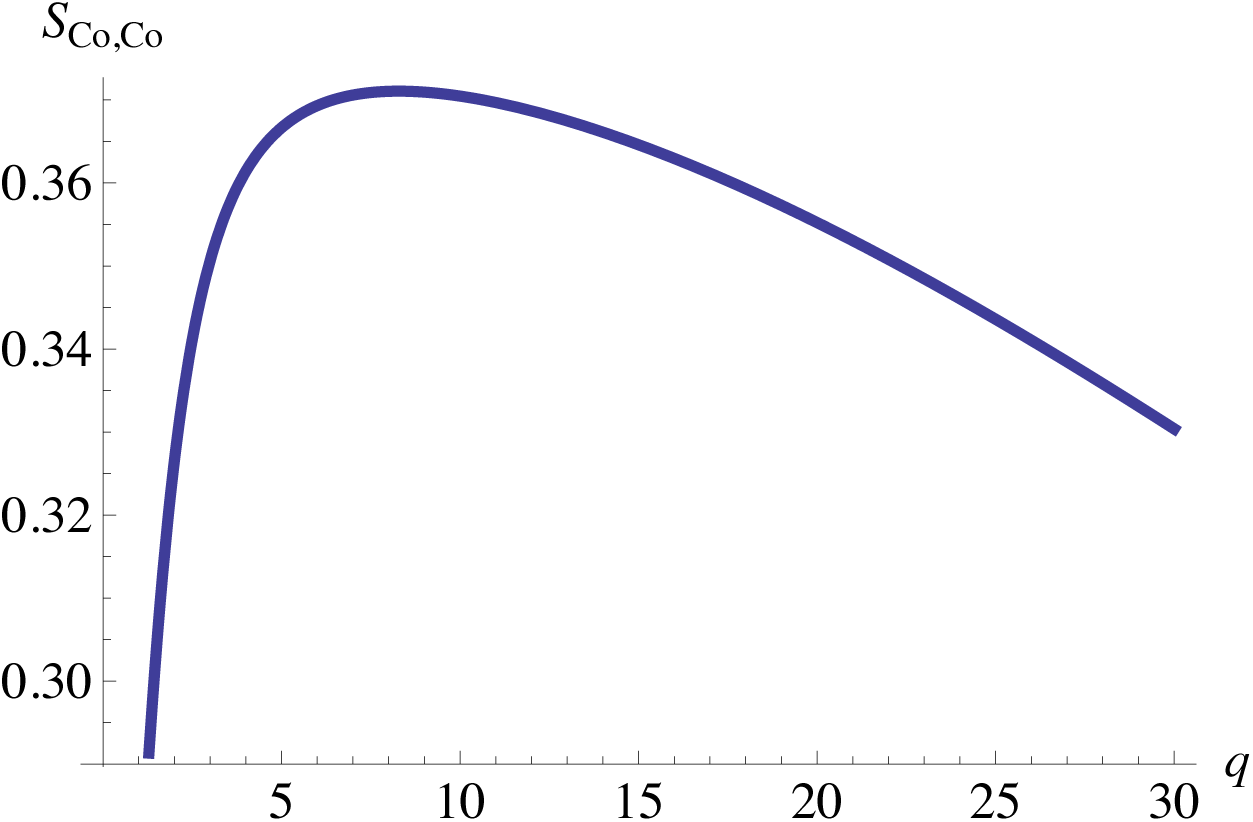
Structure function of the cholesterol in the outer leaf plotted versus the dimensionless wavevector *q* ≡ *k*(*κ*_*b*_*/σ*_*b*_)^1*/*2^. The structure function at zero *q* is *S*_*Co,Co*_(0) = 0.21.

The question now arises as to how the rafts in the inner and outer leaves are correlated. This is answered by examining the structure function of the cholesterol fluctuations in the inner and outer leaves, i.e *S*_*Co,Ci*_. This is presented in Fig. 5. As this function is negative, the regions of increased cholesterol in the two leaves are anti-correlated. By examining such structure functions, we find that the rafts consist of regions enhanced in SM and cholesterol in the outer leaf and enhanced PE and PC in the inner leaf, floating in a sea enriched in PC in the outer leaf and PS and cholesterol in the inner leaf. This configuration is illustrated schematically in Fig. 6. Due to the correlated fluctuation of cholesterol and SM, the acyl chains of the latter are expanded by the former which increases the magnitude of its negative spontaneous curvature (30). The membrane curves inward towards the cell cytoplasm and reduces its free energy. The same effect occurs with the cholesterol and PS in the inner leaf; i.e. it curves toward the outside of the cell. This can be confirmed directly from Eq. (36) that relates the fluctuations in the height to the composition fluctuations and Eqs. (15) and (16) for the spontaneous curvatures of the two leaves.

**Figure 5:**
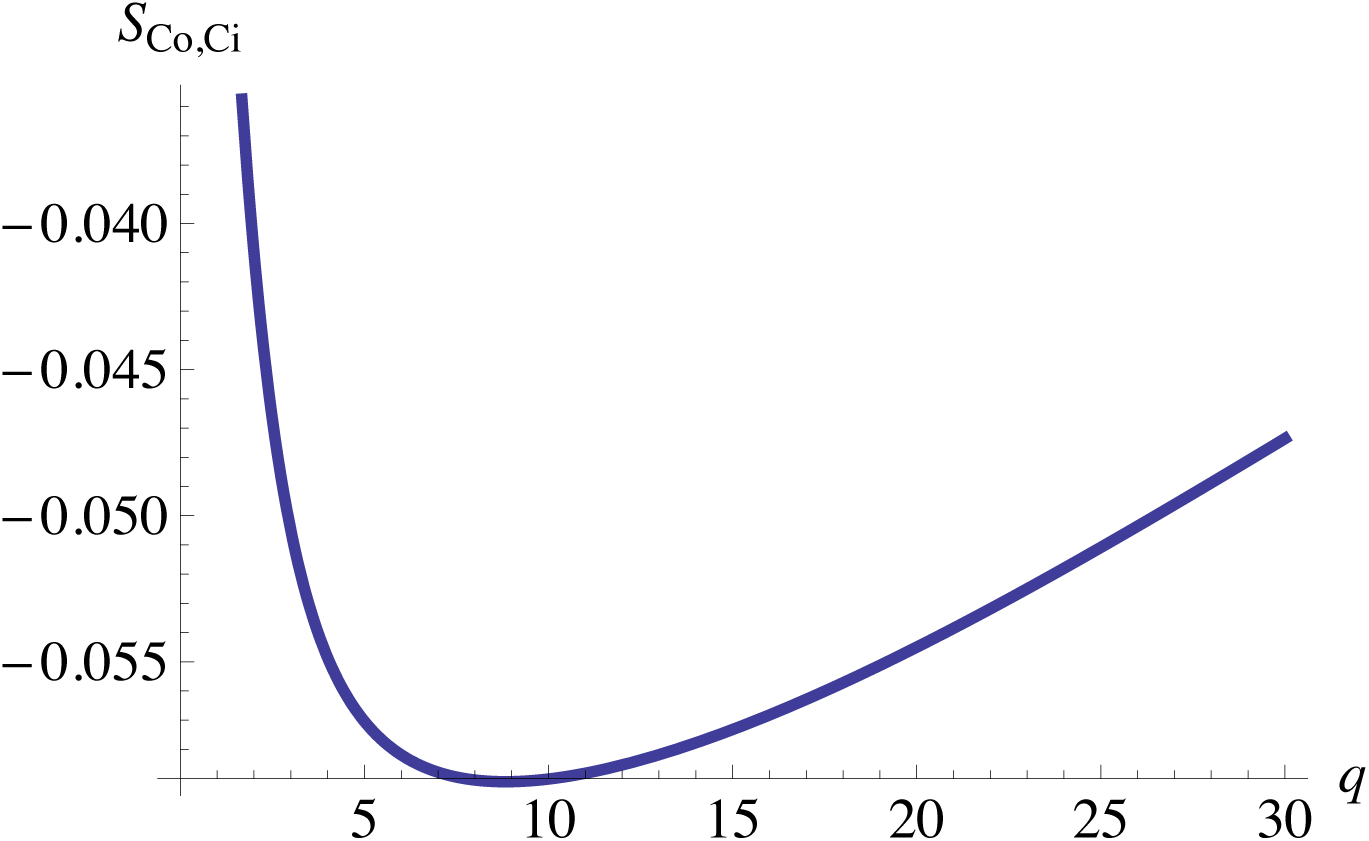
Structure function between the cholesterol in the inner and outer leaves plotted as versus the dimensionless wavevector *q* ≡ *k*(*κ*_*b*_*/σ*_*b*_)^1*/*2^. The structure function at zero wavevector is *S*_*Co,Ci*_(0) = 0.

**Figure 6:**
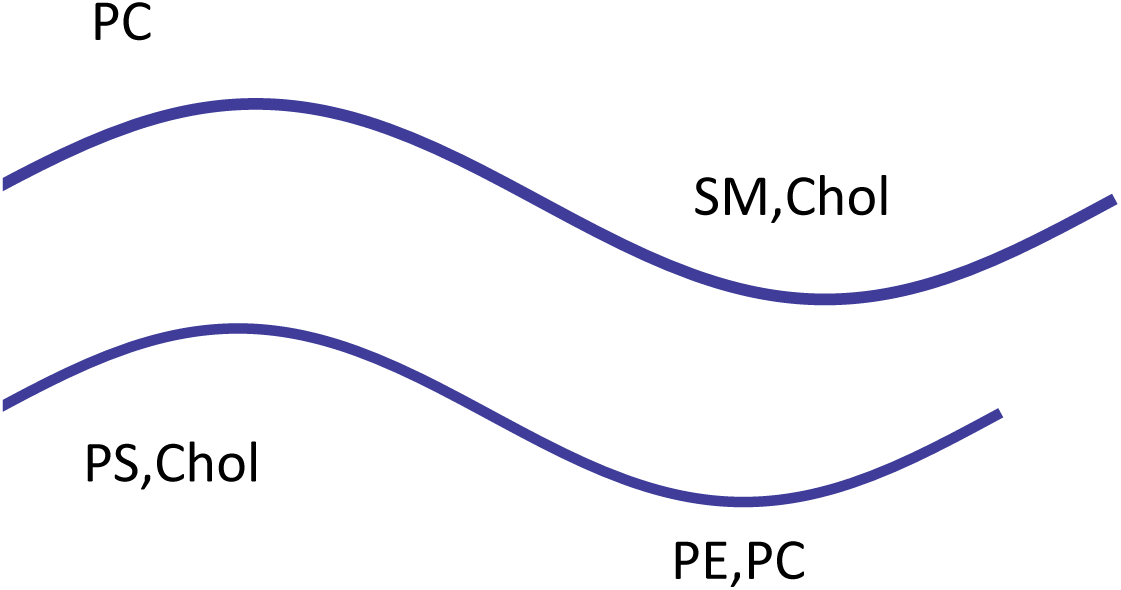
Schematic illustration of the correlated fluctuations of the components of the membrane and its spatial configuration..

Given this tendency to separate the components, we suspect that any interaction between components that has the same effect would cause the microemulsion to be stronger as evidenced in the structure factor. To test this assumption, we increase the interaction between SM and PC, which had been set to zero, as taken from Refs.(40, 41), to a value of *ϵ*_*SM,P C*_ = 0.3*k*_*B*_*T*. We do this on the basis of experimental results on the ternary mixture of SM, POPC, and cholesterol (5) that show phase separation at 37 ^*°*^C. The data require a non-zero SM-PC interaction to fit the observed binodals. The results for the structure function *S*_*Co,Co*_ are shown in Fig. 7, and should be compared with those in Fig. 4 with zero interaction between the SM and PC. In the system with the non-zero SM, PC interaction, the height of the peak in *S*_*Co,Co*_ is now more than two and one half times larger than its value, 0.33, at zero wavevector.

**Figure 7:**
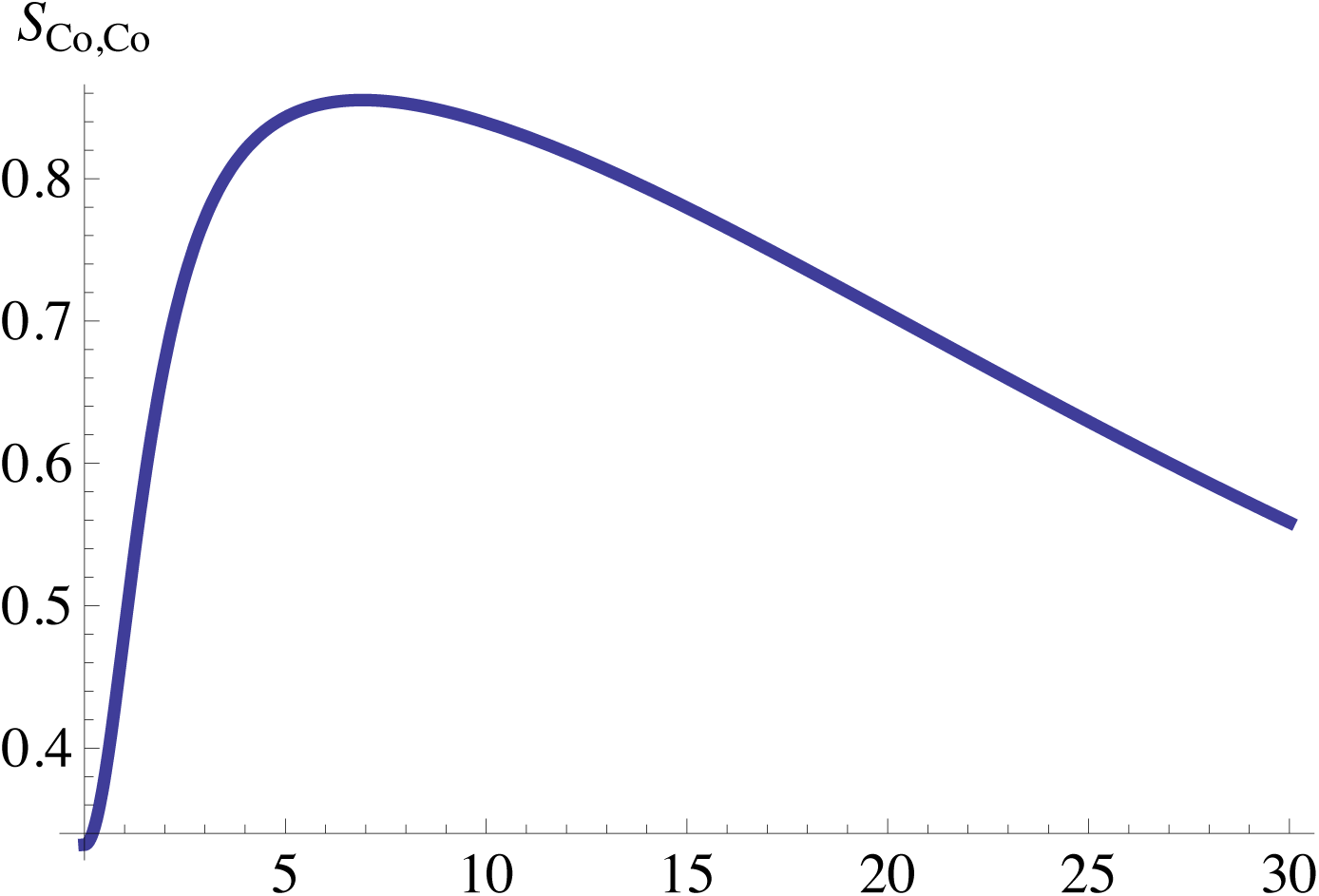
Structure function between the cholesterol in the outer leaf and with an interaction between the SM and PC of 0.3*k*_*B*_*T*. It is plotted as a function of the dimensionless wavevector *q* ≡ *k*(*κ*_*b*_*/σ*_*b*_)^1*/*2^. The structure function at zero *q* is *S*_*Co,Co*_(0) = 0.33.

One can take the Fourier transforms of the structure functions to obtain real-space correlation functions. These are not very useful, however, because the structure functions result from averages, both over the directions of the wavevector, and over the ensemble of states. Although the microemulsion contains internal structure, it still is a disordered fluid. Consequently most of the evidence of its structure in real space is lost in the averaging. More informative about the internal structure are “snapshots” from simulations, such as those in Ref. (28) from a much simpler Landau model than that investigated here. A large-scale simulation of the plasma membrane has been carried out (42) but, because it did not include height fluctuations of the membrane, no microemulsion brought about by the mechanism discussed in this paper would have been produced.

## 4. Discussion

We have employed a recent model (30) of the plasma membrane that has three components in the exoplasmic leaf and four in the cytoplasmic leaf, and calculated its free energy up to Gaussian fluctuations. Composition fluctuations are coupled to height fluctuations of the membrane. The principal result of this work is that both leaves of the plasma membrane are in a disordered liquid, microemulsion phase; that is, a liquid phase with fluctuating internal structure characterized by a well-defined length. The theory employed neatly addresses the two major problems encountered in other theories. First is the existence of a characteristic length of the internal structure, (*κ*_*b*_*/σ*_*b*_)^1*/*2^, that arises from the coupling of concentration fluctuations to height fluctuations of the membrane. As noted above, with *κ*_*b*_ = 44*k*_*B*_*T* (34) and *σ*_*b*_ = 0.00467*k*_*B*_*T/nm*^2^ (23), this length is 97 nm. Second is the question of how a raft arises in both leaves of the membrane. Within this theory, wherever the membrane is relatively of the same thickness, the mechanism couples the leaves, and the microemulsion appears in both of them. The rafts and the sea are distinguished in both leaves by their difference in curvature. In addition, because of the presence of SM in the outer leaf, the raft there is also distinguished from the sea by a difference in phospholipid areal density.

Because the height of the peak in the structure factor, which occurs at non-zero wavevector, is large compared to its value at zero wavevector, it appears that the microemulsion is a fairly strong one; with a weak microemulsion being one close to the line at which the maximum just emerges at a non-zero wavelength, the Lifshitz line, and a stronger one being closer to the transition to a modulated phase (43). The mechanism tends to separate the components into two groups; a “raft” of SM and cholesterol in the outer leaf and PE and PC in the inner leaf, and a “sea” of PC in the outer leaf and PS and cholesterol in the inner leaf. We have noted that any interactions which also promote such a separation would make the microemulsion stronger.

The underlying theory is, of course, an experimentally testable one. Indeed some of its predictions have been confirmed, such as the behavior of the characteristic domain size with temperature and surface tension. However whereas domains in a symmetric membrane would, by this theory that has no direct coupling between leaves (44, 45) be predicted to be anti-correlated, domains were reported to be corrrelated(26). There is clearly a need for additional experimental tests. In sum, we have applied the theory to a model of the plasma membrane with reasonable compositions and elastic properties, and have shown that, at biological temperatures, it exhibits a microemulsion in both leaves, and thereby provides a rational basis for the concept of “rafts”.

## 5. Author Contributions

All authors contributed equally to this work.

## 6. Acknowledgments

One of us, (M.S.) acknowledges rewarding comversations with S. Holcomb.

